# Zooplankters in an oligotrophic ocean: contrasts in the niches of the foraminifers *Globigerinoides ruber* and *Trilobatus sacculifer* in the tropical South Pacific

**DOI:** 10.1101/2019.12.31.892208

**Authors:** George H. Scott

## Abstract

The distributions of two morphologically similar planktonic foraminifera (*Globigerinoides ruber* and *Trilobus sacculifer*) that are major taxa in the mixed layer of the tropical South Pacific Ocean are related to environmental variables (sea surface temperature, chlorophyll-a, nitrate, phosphate, salinity, oxygen) to determine the extent to which their niches overlap. Their distributions in ForCenS, a database of species in seafloor sediment are studied as a proxy for upper ocean data and are analysed as occurrences using MaxEnt, and as relative abundances via non-parametric regression (Random Forests). Their distributions are similar and their co-occurrences are high but relations between their abundances and the environmental variables are complex and non-linear. In the occurrence analysis sea surface temperature is the strongest predictor of niche suitability, followed by chlorophyll-a; environments between 0 – 20° S are mapped as the most suitable for both species. To the contrary, predicted species distributions are strongly differentiated by the abundance analysis. Nitrate and chlorophyll-a are primary variables in the map of predicted relative abundances of *Globigerinoides ruber*, with maxima in the hyper-oligotrophic zone of the subtropical gyre. In contrast, sea surface temperature and chlorophyll-a are primary variables in the map for *Trilobatus sacculifer* and predicted maxima are at the margins of the hyper-oligotrophic zone and near the West Pacific Warm Pool. The high relative abundance of *Globigerinoides ruber* in the hyper-oligotrophic zone is attributed to its close photosymbiotic relation with on-board dinoflagellates; this compensates for the low primary productivity in the zone. It is clearly identified as the best-adapted planktonic foraminifer in this huge marine ‘desert’ and might serve as a useful proxy. The photosymbiotic relation is less apparent in *Trilobatus sacculifer* which, as *in vitro* research suggests, primarily depends on particulate nutrition. The study shows the value of abundance over occurrence data for analysing the trophic resources of these zooplankters.

## Introduction

Although planktonic foraminifera, with only c. 48 morphospecies (Schiebel and Hemleben, 2017) are a minor component of the zooplankton, they are particularly valuable in (paleo-) oceanography because their calcitic shells, which contain a variety of environmental and geochemical signatures (Rippert et al., 2016), are widely preserved in the bottom sediments of the world ocean. In these and other ways, including their value as a carbon sink (Schiebel et al., 2002), their small presence in the plankton belies their utility, for which an understanding of their distribution and ecology is essential. Globally, Bé (1977) comprehensively reviewed their distribution in the modern ocean and Bé & Hutson (1977), in a pioneering example of species distribution modelling (SDM) in planktonic foraminifera, estimated the optimum values of sea surface temperature (SST), salinity and phosphate for species in the Indian Ocean. More recently, modelling of species distributions (e.g., Žarić et al. 2006; Fraile et al., 2008; Lombard et al., 2011; Waterson et al., 2017; Kretschmer et al., 2018) has been advanced by improvements in hydrological databases, *in vitro* research and modelling software. These studies have been directed primarily to predicting proxy paleoenvironmental distributions, including seasonal variations in flux. Little attention has been directed to how co-occurring species in the modern ocean share environmental resources. Investigated here are the distributions of *Globigerinoides ruber* and *Trilobatus sacculifer* in the South Pacific Ocean (SPO). One basin is studied to avoid inter-basin difference in hydrology; moreover, the SPO subtropical gyre contains the world’s largest hyper-oligotrophic water mass (Morel et al., 2010). The targeted taxa are major components of foraminiferal assemblages in equatorial through warm subtropical water. They are morphologically similar, live in the mixed layer and commonly co-occur. Both are spinose, develop supplementary apertures in later ontogeny (Fig. 1 A-E) and are hosts to symbiotic algae yet *Globigerinoides ruber* is commonly more abundant than *Trilobatus sacculifer* and often dominates assemblages. These attributes suggest that the taxa might be close competitors. To what extent do their niches overlap?

**Figure 1.**
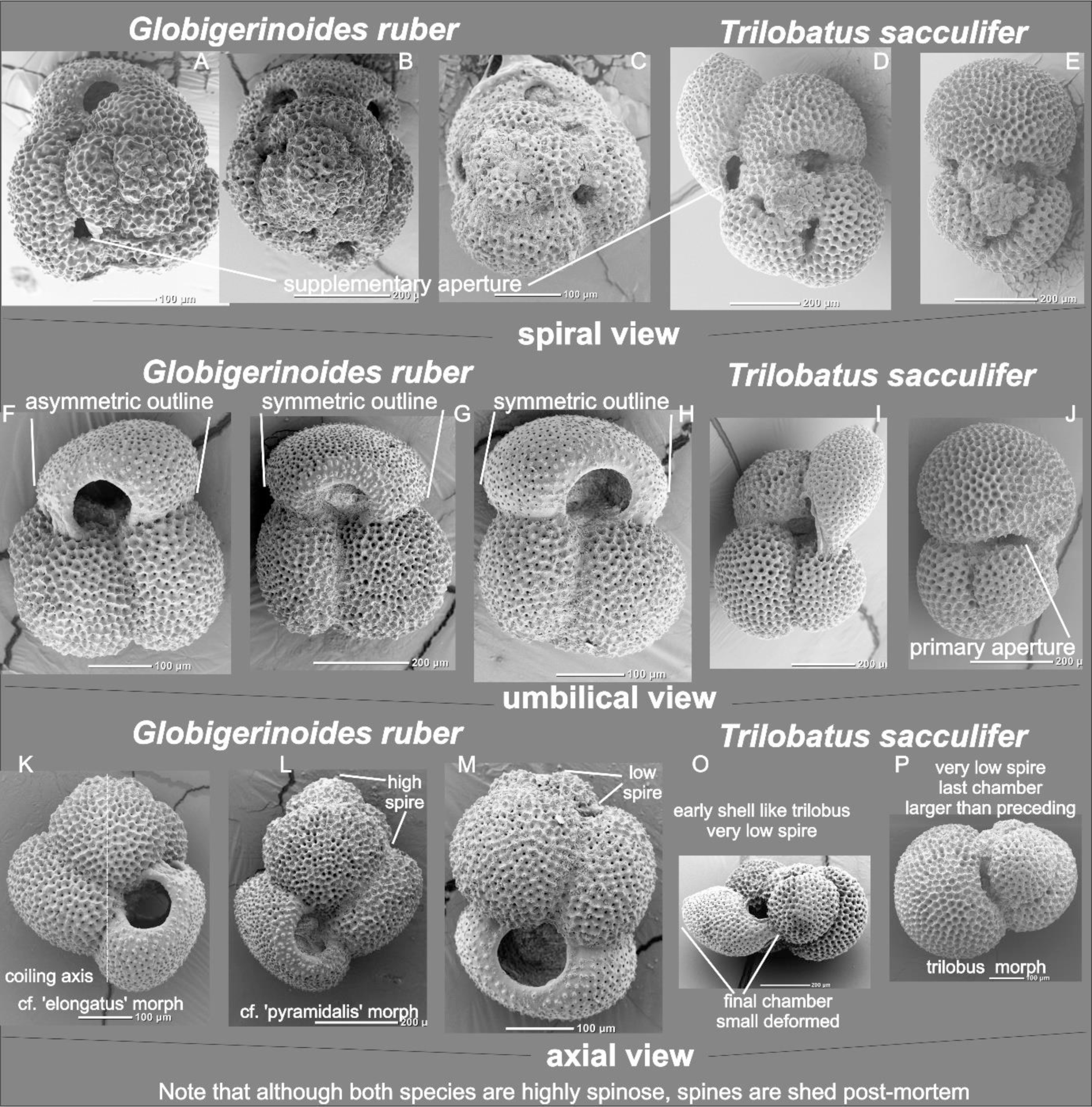
A F K, B G L, C H M spiral, umbilical and axial views of three specimens of *Globigerinoides ruber*. D I O, E J P spiral, umbilical and axial views of two specimens of *Trilobatus sacculifer*. All specimens from DSDP Site 591A core 1, section 1, 0-1 cm 31° S 164° W, Lord Howe Rise, Southwest Pacific.

Ideally, answers to this question should be founded on studies of living populations as recorded in plankton nets or traps. However, such data are sparse compared to the record available in ForCenS, a database of census records of planktonic foraminifera in bottom sediments (Siccha and Kucera, 2017). This proxy for upper ocean populations is used to model the spatial distribution of the taxa in relation to physical and geochemical properties of the upper ocean (SST, chlorophyll-a, nitrate, phosphate, oxygen, salinity). Species distribution models are extensively used in ecology for studies of niche resolution and the widespread availability of data on species occurrences has led to them becoming the principal data for building SDMs, with MaxEnt (Phillips et al., 2017) as the default tool. This study compares the suitability of this approach with niche discrimination via regression of species abundances on environmental variables. Species responses to the SPO hyper-oligotrophic zone are considered and features of the database and the integrity of the taxa as morphospecies are reviewed.

## Material and Methods

Initiation of the Deep-Sea Drilling Program and the perceived value of planktonic foraminifera to estimate paleo-sea surface temperatures (CLIMAP Project Members, 1976) led to the compilation of a substantial database of species abundances in core tops from the world ocean. Siccha and Kucera (2017) detailed the methodology used to assemble the current version, ForCenS. This database was downloaded from https://doi.pangaea.de/10.1594/PANGAEA.873570 in October 2018; relative abundances of columns labelled ‘Globigerinoides_white’ (*Globigerinoides ruber* in this study) and ‘Trilobatus_sacculifer’ (*Trilobus sacculifer* in this study) were extracted. A subset that includes records from the South Pacific Ocean (SPO) and its extension into the Southern Ocean (140°-290° E, 1° N-70° S) was used. Collection cleaning (Gomes et al., 2018) was not applied. Although nomenclature in ForCenS is carefully harmonized, errors in identification of taxa by its many contributors are unknown, as are errors in specimen counts. Samples, which are variously time-averaged, may also be affected by advection and dissolution. Species fluxes cannot be estimated from the specimen counts. In these aspects the database is a noisy proxy relative to data from sediment traps and plankton tows (e.g., Žarić et al., 2006). This study is restricted to a single basin to avoid inter-basin oceanographic effects on species distributions. Of particular interest is that the SPO contains a very large hyper-oligotrophic zone (D’Hondt et al., 2009; Morel et al., 2010) that is a major influence on plankton ecology (Williamson and McGowan, 2009).

A major advance in planktonic foraminiferal biogeography has been the introduction of geospatial techniques to model the relation between species distributions and their environmental resources (Žarić et al., 2006; Fraile et al., 2008; Lombard et al., 2011). Oceanographic variables commonly used in such studies include SST and chlorophyll-a. Here mean annual SST is from CARS 2009 (Ridgway et al., 2009) and chlorophyll-a from WCMC-034 (mean annual sea surface chlorophyll-a concentration 2009-2013, available from http://www.unep-wcmc.org/). Additionally, nitrate (umol/l), phosphate (umol/l), salinity (psu) and oxygen (ml/l) data are from CARS 2009. A comprehensive database of particulate organic matter was not available. All data were clipped to the spatial dimensions of the ForCenS subset and values obtained using inverse distance bilinear interpolation.

Because species occurrence data are widely available, MaxEnt (Elith and Leathwick, 2009; Merow et al., 2013; Phillips, 2017) is widely used in ecological scenarios (Robinson et al., 2017) and has become the default method for measuring the environmental suitability of locations; it uses a maximum entropy algorithm which is suitable for ecology. Given data on the occurrences of a species in an environmental variable space and background data that include all values of the environmental variable in that space, MaxEnt finds a model that best differentiates suitable occurrence locations from background locations. The version in the R package dismo (available from https://cran.r-project.org/web/packages/dismo) is used.

Random Forests (Breiman, 2001; Liaw and Wiener, 2002; Evans et al., 2011) is a non-parametric ensemble method which, by combining a large number of decision trees from bootstrap samples, finds a single model that is the best estimate of the target values (abundances) from the environmental variables. Each tree uses a random sample of the environmental variables and a random sample of the species abundances; overfitting is minimized. The implementation in ModelMap v. 3.4.0.1, available from https://cran.r-project.org/web/packages/ModelMap/index.html is used.

## Results

### Distribution

There are only minor differences in the areal distribution of the taxa as mapped by their convex hulls (Fig. 2 A, C). Both are recorded from stations between the equator and ~45° S; Most records are from the West Pacific Warm Pool (WPWP), including the Coral Sea, Fiji Basin and about the Ontong Java Rise. There are clusters about 10° - 20° S near the East Pacific Rise and near the Marquesas Archipelago. In contrast with the dense record on the western side of the basin there are few records adjacent to South America, apart from those near the Panama Basin. *Globigerinoides ruber* is recorded from 798 stations and *Trilobatus sacculifer* from 763 stations. They co-occur at 712 stations. Using the hypergeometric distribution as a null model, the probability of co-occurrence is 0.494.

**Figure 2.**
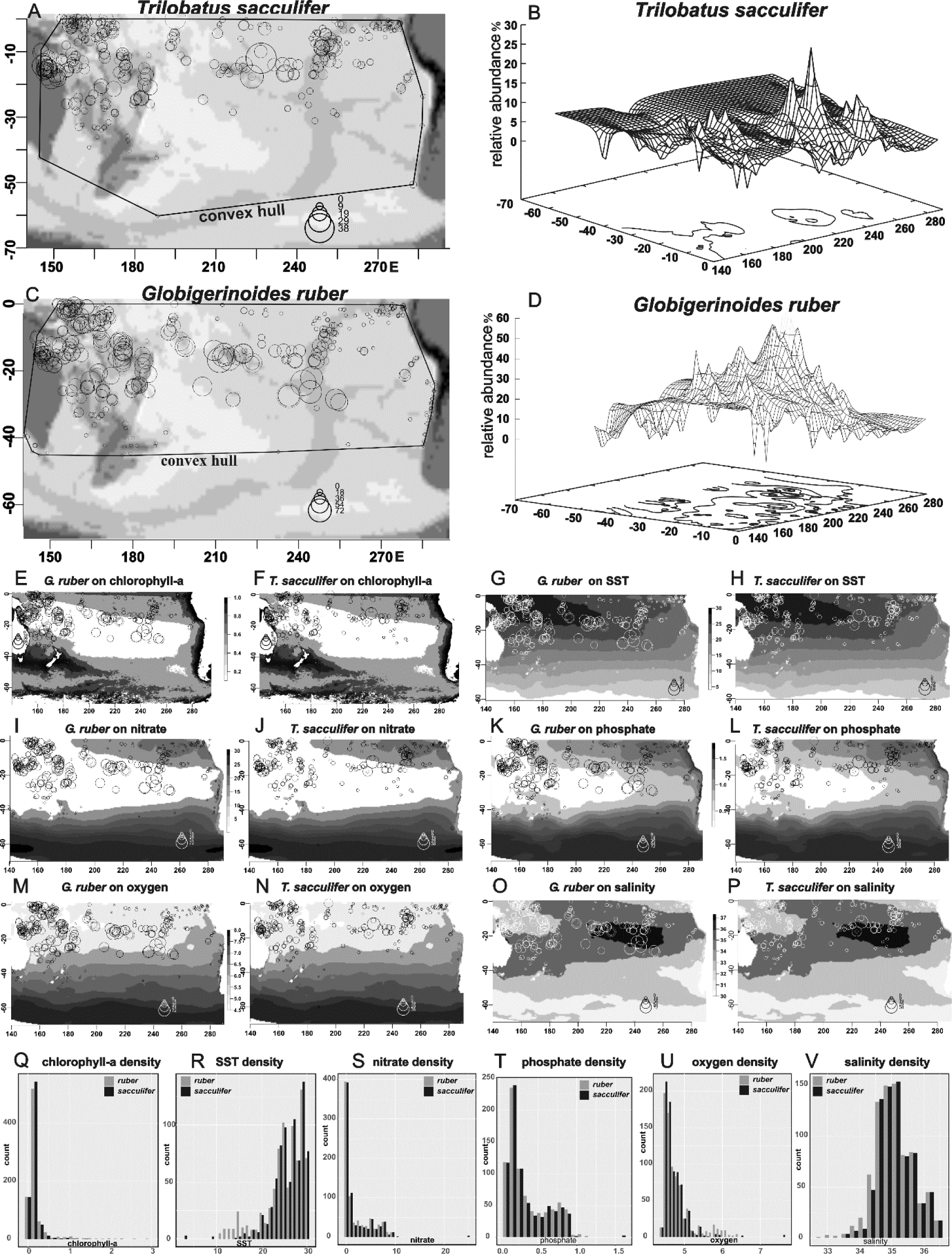
A-D 2-D and 3-D relative abundance plots of taxa in the Southwest Pacific A-B *Trilobatus sacculifer*. C-D *Globigerinoides ruber*. E-P Relative abundance of taxa plotted on distributions of environmental variables. Q-V Density histograms of environmental variables for species.

Notably, *Globigerinoides ruber* has a much larger abundance footprint in the SPO (Fig. 2 B, D) and is often the first-ranking or dominant species in subtropical assemblages, with maximum abundances of c. 70% whereas *Trilobatus sacculifer* exceeds 30% only at two stations. Abundances of the taxa in the 712 stations of co-occurrence are not significantly related in a linear regression analysis (Fig. 3 M; p < 0.01). The distribution is complex: a smooth spline fit indicates a positive relation between the taxa when they are minor members (<15%) of assemblages but the abundance of *Trilobatus sacculifer* falls near the upper tail of the distribution of *Globigerinoides ruber*.

**Figure 3.**
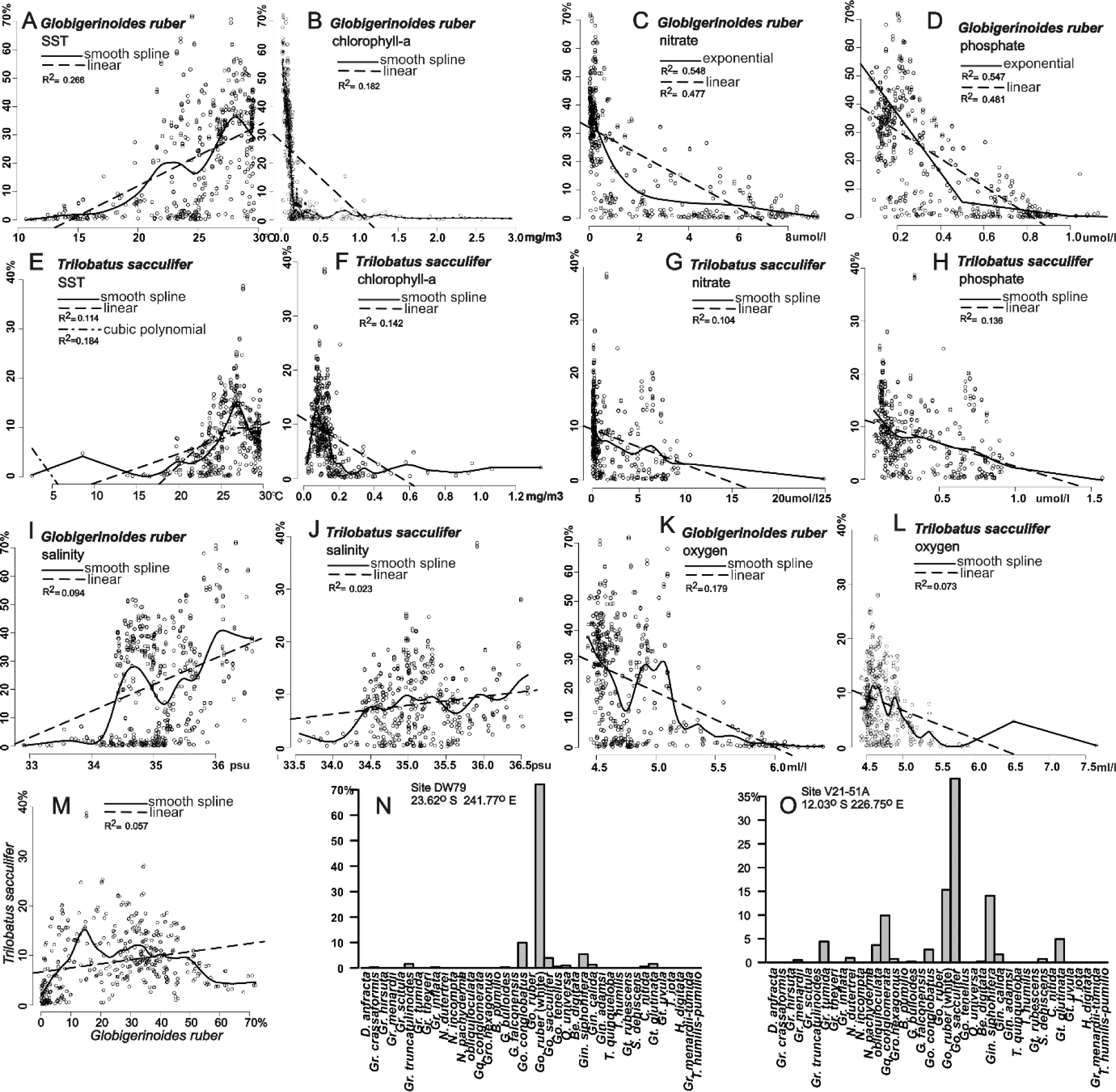
A-L Relative abundance of *Globigerinoides ruber* and *Trilobatus sacculifer* plotted against distributions of environmental variables. M Relative abundance of *Globigerinoides ruber* plotted against relative abundance of *Trilobatus sacculifer*. N Assemblage composition at a site at which *Globigerinoides ruber* is strongly dominant. O Assemblage composition at a site at which *Trilobatus sacculifer* is dominant.

### Environmental Variables

The fitted linear function between species abundance and water mass variables (Fig. 3 A-L) emphasises that relationships in all comparisons are weak and non-linear. Smoothed splines are shown to illustrate the complexity of the relationships.

The relative abundance of both species is positively related to SST but the relationship is very weak (Fig. 3 A, E); this strongly contrasts with the relation between species diversity and SST (Rutherford et al., 1999). For *Globigerinoides ruber* maximum abundances peak at ~24° C (central SPO sites) and at ~28° C (WPWP) whereas only the latter peak shows in the *Trilobus sacculifer* distribution. Apart from several questionable records of *Trilobatus sacculifer* from subantarctic stations, *Globigerinoides ruber* has a more substantial record in extra-tropical regions.

An outstanding feature in the environmental dataset is the tight association of records (*Globigerinoides ruber* 80%, *Trilobatus sacculifer* 86%) with water in which the concentration of chlorophyll-a is <0.2 mg m^−3^ (Fig. 2 E, F). Although these gross data indicate a weak, negative relation between relative abundance and chlorophyll-a (Fig. 3 B, F) the geographic distribution is well-defined, particularly for *Globigerinoides ruber* whose highest abundances (Fig. 2 E) are in the central SPO within the core of the subtropical gyre. A particular interest in this distribution is that the species is dominant in a zone of hyper-oligotrophy which has very low primary productivity and has been described as a marine desert (Raimbault et al, 2008; Morel et al., 2010; Reintjes et al., 2019). *Trilobatus sacculifer* is most abundant at sites near its northern margin in the central SPO and is common in the Coral Sea and WPWP.

Nitrate and phosphate concentrations are indirect measures of primary productivity in the oceans. Expectedly, their distributions might parallel that of chlorophyll-a. A zone of minimal nitrate concentration encompasses the minimal chlorophyll-a zone and, in the western SPO, extends to 40° S (Fig 2 I-J). The zone thus includes most extra-equatorial records of the taxa. The extent of the minimal phosphate zone (Fig. 2 K-L) is a westward contraction of the minimal nitrate zone in the central SPO but still includes the WPWP. Some of the highest relative abundances of the taxa are at stations beyond the minimal phosphate zone. *Trilobatus sacculifer* tends to be moderately common (up to 20% relative abundance) over a wider range of concentrations of these variables than does *Globigerinoides ruber*. The linear fits of the abundance of *Globigerinoides ruber* to nitrate (R^2^ = 0.477) and phosphate (R^2^ = 0.481) are the highest in the study (Fig. 3 C-D).

Salinity is highest about the hyper-oligotrophic zone in the central SPO and includes stations with high relative abundance of *Globigerinoides ruber* and *Trilobatus sacculifer* (Fig. 2 O-P). The highest abundances of *Globigerinoides ruber* are in water in which salinity exceeds 36 psu. However, that both taxa are also common in the WPWP, an extensive region of low salinity, suggests that the relation there between salinity and abundance is weak, as indicated by the linear fits (Fig 3 I-J).

Both taxa are in water with 4.5 - 5.5 ml/l oxygen (Fig. 2 M-N). *Globigerinoides ruber* (Fig. 3K) includes a tail of stations with abundances <10% that are in water with oxygen > 5.5 ml/l. The linear relation between oxygen and abundance is very weakly negative. (Fig. 3 K-L). Oxygen concentrations appear to have minor effect on the abundances of *Globigerinoides ruber* and *Trilobatus sacculifer*; this may be consistent with data in Kuroyanagi et al. (2013).

### Species Distribution Models

The objective is to find relationships between the geographic distribution of a species and its environmental resources which enable predictions about its spatial distribution (Hijmans and Elith (2017).

### Occurrence data - MaxEnt models

Sea surface temperature is the strongest predictor of environmental suitability (~70%, Fig. 4 I, K) for both taxa. Nitrate and phosphate are lesser contributors while chlorophyll-a is the lowest-ranked contributor. Predictive maps for *Globigerinoides ruber* and *Trilobatus sacculifer* are closely similar (Fig. 4 A-B), with high estimates of suitability across the equatorial SPO and particularly about the WPWP and southward from the Coral Sea along coastal Queensland, Australia. Neither model predicts that the hyper-oligotrophic central SPO is a favourable habitat. One difference is that the Humboldt Current System off South America and offshore New Zealand map as favourable for *Globigerinoides ruber* but not for *Trilobatus sacculifer*. There are high concentrations of chlorophyll-a in these regions. Nevertheless, confirming the apparent similarity of the environmental suitability maps, the Warren et al. (2008) test for niche equivalency reports that the niches of the two species are identical (p.value = 1).

**Figure 4.**
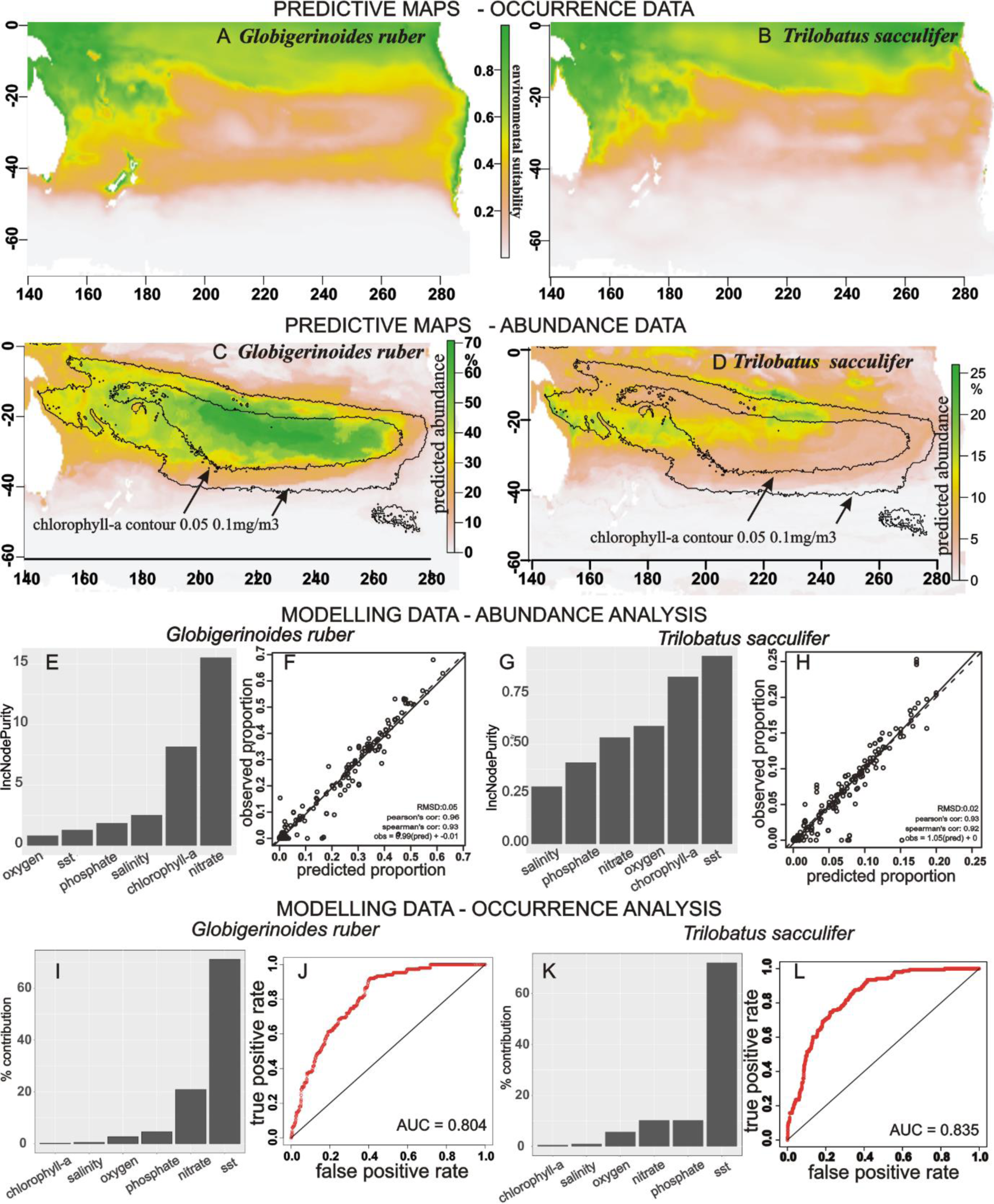
A-B Spatial distribution of environments suitable for *Globigerinoides ruber* and for *Trilobatus sacculifer* modelled from their occurrences using MaxEnt. C-D Spatial distribution of relative abundances predicted by Random Forests using abundance data. E-H Node purity, a measure of environmental variable importance, and regression statistics for the Random Forests analyses. I-L Environmental variable contributions and model fit data for the MaxEnt analyses.

### Abundance data - Random Forests models

Nitrate and chlorophyll-a are the most important predictors (Fig. 4 E) in the *Globigerinoides ruber* model; predicted and observed values of abundance are highly correlated and the model fit is good (Fig. 4 F). The predicted distribution map (Fig. 4 C) defines an extensive region in the central SPO in which *Globigerinoides ruber* is a dominant member of planktonic foraminiferal assemblages, with mean relative abundances > 45%. It is closely confined within the 0.1 mg m^−3^ chlorophyll-a contour and approximates the SPO hyper-oligotrophic zone (Raimbault et al., 2008; Morel et al. 2010).

Sea surface temperature and chlorophyll-a are the most important predictors in the *Trilobatus sacculifer* model and, as with *Globigerinoides ruber*, the model fit is good. However, in strong contrast with the occurrence analyses, their predicted distribution maps differ substantially. Highest predicted abundances of *Trilobatus sacculifer* are in the vicinity of the Tuamotu Ridge and Marquesas Islands in French Polynesia and lie beyond the 0.1 mg m^−3^ chlorophyll-a contour. Also defined is a zone of lesser abundance that extends along the path of the South Equatorial Current (Kessler and Cravatte, 2013; Ganachaud et al, 2014) from the central SPO westward to the Queensland coast.

## Discussion

### Database

ForCenS is unusual amongst natural history collections because it is a global record of planktonic foraminiferal abundances whereas most collections are regional and record only occurrences of taxa. But like most natural history collections (vanderWal et al., 2009) it is a compilation of sites that were sampled for various oceanographic and geological investigations which incidentally provided sediment suitable for planktonic foraminiferal censii. This is preferential sampling (neither random nor systematic, Pennino et al., 2019) which exposes the database to spatial bias in its coverage: some regions may be more closely sampled than others. Here it is apparent (Fig. 2 A-D) that the western tropical region of the SPO has been preferentially sampled for both species. MaxEnt is particularly sensitive to spatial bias because it assumes that the studied region has been systematically surveyed and randomly sampled (Elith et al., 2011; Phillips et al., 2017). The problem is compounded when, as in this study, the taxa are highly co-occurrent. While spatial filtering (e.g., Kramer-Schadt et al., 2013; Pennino et al., 2019) may alleviate the problem, it is not pursued here as MaxEnt is designed for occurrence data and the study clearly shows, as does Gomes et al. (2018), that abundance data are more informative.

### Occurrence vs abundance

The occurrence model predicts (Fig. 4 A-B) that the most suitable habitat for both species is in the equatorial – subtropical region of the SPO, with the WPWP the most preferred. This result is un-remarkable, biogeographically, as it accords with research that recognizes that *Globigerinoides ruber* and *Trilobatus sacculifer* are major taxa in those regions of the world ocean (Schiebel & Hemleben, 2017). Rather, the interest of the result is in the close similarly of their distribution maps and the high probability that their niches overlap. Nominally, this contravenes the competitive exclusion principle of ecological theory (Hutchinson, 1961; Sakavara et al., 2018) which asserts that such species need to differentiate their niches to survive. However, the veracity of the occurrence analysis is challenged by the analysis of the abundance data, particularly for *Globigerinoides ruber*. Its highest relative abundances are predicted to lie in the central SPO hyper-oligotrophic zone much of which, using the occurrence data, predicts low suitability for the species. Bé and Hutson (1977) noted that its abundance increased in the oligotrophic Indian Ocean Central Watermass.

To clarify this apparent paradox the trophic behaviours of the species are considered. Primarily, planktonic foraminifera are passive ambush feeders (Kiorboe, 2011): prey are trapped, not pursued. *Globigerinoides ruber* and *Trilobatus sacculifer* combine this strategy with a symbiotic relation with algae, particularly dinoflagellates. The Random Forests variable importance index (Fig. 4 E, G) and maps of predicted abundance (Fig. 4 C-D) provide clues about their use of trophic resources in the SPO. For *Globigerinoides ruber* the analysis indicates that nitrate and chlorophyll-a are primary determinants of its relative abundance. Nitrate is an input to primary producers (Chavez et al., 2011); chlorophyll-a is a measure of phytoplankton abundance, the primary producers. That the relative abundance of *Globigerinoides ruber* is greatest in a zone where values for nitrate and chlorophyll-a are notably low and prey are scarce, suggests that its trophic resources are based on a photosynthetic symbiotic relation with on-board dinoflagellates who act as local primary producers (Gast and Caron, 2001; Shaked et al., 2006). Gastrich and Bartha (1988) concluded from *in vitro* research that a symbiotic relation with algae is possibly essential for *Globigerinoides ruber* to survive in nutrient-limited environments.

Whereas nitrate is the dominant predictor variable in the abundance model for *Globigerinoides ruber*, contributions to the *Trilobatus sacculifer* model are distributed amongst the variables. Its abundance model reflects a positive relation with SST and salinity but negative with chlorophyll-a, nitrate, phosphate and oxygen. A principal effect in the predictive map is that the close relation with the minimal chlorophyll-a contour is lost (Fig. 4 D). Maximum abundances in the central SPO map northward of the hyper-oligotrophic zone, and in the Coral Sea. There is a zone of c. 10 - 20% abundance that aligns with flows in the SEC, the westward limb of the subtropical gyre. This transects the minimal chlorophyll-a contours.

At the *in vitro* level Bé et al. (1981) found that specimens of *Trilobatus sacculifer* were unable to grow if deprived of particulate food and left reliant on symbiont photosynthesis and possibly bacteria for nutrition. Takagi et al. (2018) corroborated this result. They found no direct evidence that the symbionts contributed to the nutrition of the host. Takagi et al. (2019 Table 1) reported that *Trilobatus sacculifer* hosts the same dinoflagellate *Pelagodinium béii*) as *Globigerinoides ruber* and classified the relationship for both species as obligate. An index of the species photosynthetic activity is similar (*Globigerinoides ruber F_v_/F_m_* = 0.49; *Trilobatus sacculifer F_v_/F_m_* = 0.51). On a trophic gradient from heterotrophy to permanent endosymbiosis (Takagi et al. 2019 Fig. 11) using photosynthetic variables obtained from fluorometry, *Trilobatus sacculifer* is more advanced than *Globigerinoides ruber*. However, the heterotrophic requirement of *Trilobatus sacculifer* apparently limits its populations in hyper-oligotrophic environments of the SPO except in regions where SEC eddy activity promotes biomass by nutrient uplift.

### Comparison with mechanistic models

Presented here are comparative models that relate several hydrological variables in upper ocean water masses to the distribution of *Globigerinoides ruber* and *Trilobatus sacculifer*. From a quite different basis Lombard et al. (2011) built mechanistic models (Peterson et al., 2016) of the environmental conditions needed for growth of individuals of these taxa using *in vitro* or estimated values for growth parameters (nutrition, respiration, photosynthesis). They extended the individual growth models for these species to population growth under natural conditions and to relative abundance estimates: chlorophyll-a was taken as a productivity indicator and as a measure of food availability. Comparison of modelled relative abundance with observed core top abundance in the MARGO database (a predecessor of ForCenS) gave model fits (R^2^) of 0.46 for *Globigerinoides ruber* and 0.56 for *Trilobatus sacculifer* (Lombard et al. 2011 Table 3). Their mechanistic relative abundance map for *Globorotalia ruber* (Lombard et al. 2011 Fig. 7) shows highest abundances in the SPO hyper-oligotrophic zone, comparable with that in the map for Random Forests predictions. Their map for *Trilobatus sacculifer* does not identify the hyper-oligotrophic zone as a region in which its abundance is low. Nevertheless, further mechanistic modelling with *in vitro* data should elucidate the biochemical processes that enable *Globigerinoides ruber* to better adapt to hyper-oligotrophy than *Trilobatus sacculifer*.

### Taxa

Underlying the integrity of their records in ForCenS and in most studies of living populations are the concepts of *Globigerinoides ruber* and *Trilobatus sacculifer* as morphospecies. These are almost wholly based on qualitative perceptions of shell architecture from optical imagery. The observed wide variability in the architecture of planktonic foraminifera, particularly evident in late ontogeny, has been simplified and structured by the proposal of morphospecies, usually using holotypes as exemplars (Scott, 2011).

Poole and Wade (2019) interpreted *Trilobatus sacculifer* (Fig. 1 D-E, I-J, O-P) as a plexus of morphospecies (*Globigerina quadrilobata* (d’Orbigny, 1846); *Globigerina triloba* (Reuss, 1850); *Globigerina sacculifera* (Brady, 1877); *Globigerinoides sacculifera* var. *immatura* (LeRoy, 1939)) in the modern ocean. Although they accepted molecular data (André et al., 2013) that the plexus probably represented a single species, as it is treated in ForCenS and Schiebel and Hemleben (2017), they considered that all were valid taxa because of their value in biostratigraphic and paleoecologic research.

A wide spectrum of shell forms has also been identified as *Globigerinoides ruber* (Fig. 1 A-C, F-H, K-M): their interpretation is complicated by evidence of multiple genotypes (Aurahs et al., 2009; 2011). In the modern SPO Parker (1962) recognized three morphogroups based on perceived and measured differences in shape and size of chambers, shell, and apertures. She viewed them as latitudinal phenotypic variants but noted that one group resembled *Globigerina elongata* (d’Orbigny 1826); cf. Fig. 1 A, F, K. Aurahs et al. (2011) produced molecular and morphometric data which showed that the Type 11a genotype represented this morphospecies. André et al. (2014) revised the molecular data and showed the presence of the Type 11a genotype in the WPWP, accompanied by several other genotypes. While molecular data identify the presence of several genotypes in the SPO, recognition of their morphological proxies is impeded by the paucity of morphometric data on the size and shape of chambers formed in latest ontogeny. Possibly some of the morphotypes in the North Atlantic subtropical gyre, where it is the dominant species (Bonfardeci et al., 2018), are present as intrapopulation variants. Here *Globigerinoides ruber*, as classified as ‘*Globigerinoides*_white’ in ForCenS, is regarded as an omnibus taxon concept of uncertain taxonomic integrity.

## Conclusions

As a natural history database of counts of planktonic foraminiferal taxa in sediment samples ForCenS is a useful proxy for their distribution in the upper ocean but its sampling basis is preferential. This impacts particularly on its use for analyses of species occurrence data. High co-occurrences of records of *Globigerinoides ruber* and *Trilobatus sacculifer* in the SPO leads to replicate sampling of oceanographic variables (Fig. 2 Q-V) and to recognition that complete niche overlap is highly probable. Analysis of abundance data which, as Gomes et al. (2018) demonstrated, is often significantly more informative, does not support this result.

Species abundances are closely predicted by non-parametric regression (Random Forests algorithm) using the same set of primary hydrological variables. These analyses show that *Globigerinoides ruber* and *Trilobatus sacculifer* respond differently to variation in some major variables. *Globigerinoides ruber* appears to survive in the impoverished hyper-oligotrophic environment of the central SPO subtropical gyre by utilization of nutrients from on-board symbionts, effectively overriding their paucity in the water mass. However, as the data are relative abundances, its dominance of planktonic foraminiferal assemblages does not necessarily imply that it flourishes in hyper-oligotrophic environments. Rather, it identifies that it is the most successful species in a highly inhospitable environment. High relative abundances (~60%>) may serve as a useful proxy for hyper-oligotrophic water masses. *Trilobatus sacculifer* appears to lack this photosymbiont faculty and its abundance distribution likely reflects its dependence on particulate organic matter. While the geographic distribution of these species overlap, their trophic niches differ.

## Acknowledgments

I am most grateful for comments on an earlier draft from Martin Crundwell and Joe Prebble (GNS Science) and from Haruka Takagi (University of Tokyo).

